# Maternal separation induces autism spectrum disorder in young rhesus monkeys

**DOI:** 10.1101/2022.03.17.484827

**Authors:** Xiao-Feng Ren, Shi-Hao Wu, Hui Zhou, Long-Bao Lv, Zi-Long Qiu, Xiao-Li Feng, Xin-Tian Hu

## Abstract

Autism spectrum disorder (ASD) is a class of severe neurodevelopmental disorders with a high incidence in young children, and its pathogenesis remains elusive. There is no effective treatment, and ASD children usually have a hard time in integrating into society and leading a normal life, which places a heavy burden on the families and society. Studies have shown that in addition to genetic factors, environmental factors are another important risk contributing to the pathogenesis of ASD. Early environmental adversity, which can lead to abnormal brain development and affect cognition and behavior, greatly increases the incidence of a variety of brain developmental diseases including ASD. However, studies on this aspect are inadequate at present, and no clear conclusions can be drawn. We explored whether early adversity could trigger ASD core clinical symptoms in macaques by modeling early adversity through maternal separation. In this study, we conducted a rigorous behavioral analysis of 12 male macaques (1.5-2 years old) that underwent maternal separation and 9 male normal macaques of the same age that had been mother raised, and found that maternal separation could induce a small number of the young individuals to develop three core symptoms of ASD, including social impairment, stereotyped behaviors, and restricted interest simultaneously. According to DSM-V and ASD clinical diagnostic criteria, these individuals should be ASD macaques for having all the three core ASD symptoms at the same time., For the first time, this study reveals that early environmental adversity can lead to ASD pathogenesis in monkeys, and provides a new approach for future ASD studies and modeling ASD monkeys.

## 1. Introduction

Autism spectrum disorders (ASD) is a type of brain development disorder that occurs in infancy and seriously affects children’s brain development and mental health (Maiano, Normand et al. 2016, Hodges, Fealko et al. 2020). The classic clinical symptoms are impairments in social interaction, restricted interest, as well as repetitive, stereotyped behaviors (Diagnostic and Statistical Manual of Mental Disorders-5th Edition, DSM-V) (Zhao, Jiang et al. 2018). ASD can cause serious damages to children’s mental health, and 60%-70% of ASD patients are not able to participate in normal education and work, which usually lead to lose the ability to live independently and cause a heavy burden on the family and society (Hodges, Fealko et al. 2020).

Epidemiological studies have shown a significant increase in the incidence of ASD in recent years (DeFrancesco 2001, Previc 2007). According to data released by US Centers for Disease Control and Prevention (CDC) in 2021, ASD prevalence was as high as 2.27%, a 23% increase from two years ago, and 3.39 times more than the prevalence first reported by the CDC in 2000 (Maenner, Shaw et al. 2021). However, as ASD is caused by multiple factors, its etiology is very complicated, and its pathogenesis is unclear. There is no effective treatment at present.

Studies have shown that both genetic and environmental factors are ASD pathogenic factors (De Rubeis, He et al. 2014, Bourgeron 2015, Krumm, Turner et al. 2015, Lappe 2016, Karimi, Kamali et al. 2017). Sufficient data have shown that heredity is one of the important risk factors of ASD (Ghosh, Michalon et al. 2013). Many gene mutations such as neuroligin 3/4, Shank3, neurexin 1, MeCP2, Hoxa1, PTEN, UBE3A and MEF2C have been found to be closely linked to the pathogenesis of ASD (Lintas and Persico 2009, Qiu and Cheng 2010). In recent years, with the rapid advance of gene editing technology, the research on the pathogenic genes of ASD has made significant progress. In particular, several scientists’ groups have made significant contributions to the development of transgenic ASD monkey models (Jennings, Landman et al. 2016). In 2016, Qiu Zilong’s group established the first lentiviral MeCP2 transgenic cynomolgus monkey model of autism (Liu, Li et al. 2016). In 2017, our group and Dr. Ji Weizhi’s group used TALEN technology to knock out MeCP2 gene, which made cynomolgus monkey exhibit core ASD symptoms such as stereotyped behaviors and impaired social interaction together (Chen, Yu et al. 2017). In 2019, Feng Guoping’s group edited the Shank3 gene of macaques, which induced the phenotype of ASD in macaques (Zhou, Sharma et al. 2019). In 2021, our group and Qiu Zilong’s group used in situ gene editing technology to edit the MeCP2 gene in the hippocampus of young macaques, and successfully induced the classic phenotypes of ASD in young macaques (Wu, Li et al. 2021).

Although most current ASD studies have focused on ASD related genes (Niu, Shen et al. 2014, Liu, Li et al. 2016, Yao, Liu et al. 2018, Kang, Chu et al. 2019), research over the past 20 years has also revealed that environmental factors, particularly those in early life, are another important risk factor of ASD (Courchesne 2002, Lappe 2016, Karimi, Kamali et al. 2017), because early life is a critical and sensitive period of brain development. Adverse early environment would lead to abnormal brain development, which in turn affects cognition and behavior, and greatly increases the incidence of many developmental brain diseases, including ASD (Binder 2017).

In 2011, Hallmayer et al. analyzed 192 pairs of human twins with at least one ASD patient, and assessed the risk of ASD genetic factors. The results showed that the shared environment of twins may play a more important role in the development of ASD than genes. The environmental factors of twins explained about 55% of ASD risk, and genetic factors less than 40% (Hallmayer, Cleveland et al. 2011). In 2014, Sandin et al. analyzed data of all 2 million children born in Sweden from 1982 to 2006, of which 14,516 were diagnosed as ASD. This is the largest study to date assessed the proportion of ASD risk that may be due to genetic factors. The results showed that for the pathogenesis of ASD, the importance of environmental factors is basically the same as that of genes. The heritability of ASD is 50%, and the other 50% are explained as non-genetic or environmental factors. Based on these work and other findings, research on early environmental risk factors for ASD has gradually gained attention (Sandin, Lichtenstein et al. 2014).

In humans, early adversity refers to life experiences that occur during infancy and childhood and cause physical or psychological harm to the individual, such as lack of parental companionship, parental divorce, abuse, witnessed parental violence, or experiencing a major natural disaster (Dahl, Larsen et al. 2017, Opendak, Gould et al. 2017). Numerous studies have shown that early adversity is an important negative environmental factor, which can alter neuronal plasticity and brain development at early age, resulting in long-term and profound negative impact on the brain (Binder 2017). Several recent studies have shown that rats that the brain plasticity of rats who have experienced maternal separation (a commonly used method to mimic early life adversity) was abnormal, and could induce ASD-like symptoms such as stereotyped behaviors and social impairment (Bahi 2017, Kaidbey, Ranger et al. 2019, Mansouri, Pouretemad et al. 2020). However, there are few similar studies on humans and non-human primates. Whether early adversity after birth could induce ASD is not clear.

Early adversity in humans is often simulated by animal models of maternal separation. The currently widely used rodent models are very different from humans in brain structure and function, behaviors and cognition (Jennings, Landman et al. 2016). The complex symptoms of human diseases are often difficult to be faithfully replicated in rodents. For example, the effects of maternal separation on humans and rodents are different: the negative effects of maternal separation on human brains and behaviors are long-term and usually cannot be reversed by later normal social life. Whereas in rodents, the negative effects of maternal separation can be reversed by a subsequent normal social life (Feng, Wang et al. 2011, Binder 2017).

Because monkeys are highly similar to humans in terms of gene background, physiological structures, brain structures and functions, human disease models developed by monkeys are better in terms of replicating the disease symptoms and simulating disease mechanisms and pathological processes (Qin, Chu et al. 2015, Li, Su et al. 2021, Li, Yao et al. 2021). For example, one of our previous studies demonstrated that the negative effects of maternal separation on monkeys are also long-term and cannot be easily reversed by subsequent normal social life (Feng, Wang et al. 2011). In other words, the macaque model of maternal separation is able to simulate the early adversity in humans faithfully. Moreover, macaques have high cognition, complex social activities and sophisticated motor control, which are particularly important for effective evaluation of the core symptoms of ASD, such as social interaction impairment, stereotyped behaviors and restricted interest, both are based on fine motor control.

Present study was designed to explore whether maternal separation could induce ASD monkeys, which should have all the core clinical ASD symptoms: impaired social interaction, stereotyped behaviors and restricted interest simultaneously. We initially selected monkeys with at least one ASD core symptoms from 12 male young monkeys having undergone maternal separation. Then, each selected candidate monkey underwent a more detailed behavioral analysis to examine whether it exhibited all three ASD core symptoms simultaneously to determine it was an ASD monkey.

## 2. Methods

### 2.1 Experimental animals and ethics

The twenty-one experimental animals for this study were healthy young rhesus monkeys (*Macaca mulatta*) (male, 1-1.5 years old, 1.5-2.5 kg) purchased from Kunming Primate Research Center, Chinese Academy of Sciences. All monkeys were treated in accordance with the National Institute of Health (USA) Guide for the Care and Use of Laboratory Animals. All animals had free access to water and were fed with monkey chows supplemented with seasonal fresh fruit and vegetables twice a day. The experimental protocol had been approved by the Ethics Committee of Kunming Primate Research Center (AAALAC accredited, IACUC16004) affiliated to Kunming Institute of Zoology, Chinese Academy of Sciences.

The twenty-one male macaques in this study were divided into maternal rearing group (MR group, n = 9) and maternal separation group (MS group, n = 12). The infant monkeys of the MR group were reared by their mothers for the first 3 months after birth and lived in several reproductive colonies of the same layout: an indoor room (2.61×2.46×2.58 m) connected to an outdoor big cage (2.67×2.66×2.67 m). Each colony consisted of 1 adult male macaque and 4-5 adult female macaques and their offsprings.

The infant monkeys of the MS group were separated from their mothers at birth for following reasons. First, about 60% of maternal separation was due to the inexperience of first-time monkey mothers. Some of them got panic and did not know how to handle the infant monkeys properly, which would cause serious injury or even death to the monkey babies. In this case, the breeders would take the monkey babies and keep them in a constant temperature incubator to increase the survival chance of newborn monkeys. Second, nearly 20% of infant monkeys were forced to be separated from their mothers because the mothers did not have adequate milk. Third, about 20% of maternal separation was due to rainy, cold weather. Born in cold weather would cause the newborn monkeys to be wet and hypothermic, which could lead to illness or even death. To protect them, the breeder would temporarily take them away from their mothers and feed them in constant temperature incubators. When the weather got better, they were brought back to their mothers. A subset of them refused to take back their babies, who then would be artificially raised in the incubators.

The infant monkeys of the MS group were fed in the incubators by professional breeders for the first month. The temperature in the incubator was maintained at 32 ± 1 °C with a dry towel on the bottom. The breeder fed the infant monkey milk 7-9 times per day, changed the towel and cleaned the incubator regularly. When the infant monkeys reached 1 month of age, they were transferred to indoor cages (0.74×0.71×0.74 m) and raised in pairs until 3 months of age. During this period, the room temperature was maintained at 21 ± 2 °C and the relative humidity was about 60%, with a normal light / dark cycle (14h/ 10 h, the lighting period was from 7:00 to 21:00).

When the monkeys reached 6 months of age, they were grouped into several social groups living in colonies of the same layout: an indoor room (2.61×2.46×2.58 m) connected to an outdoor big cage (2.67×2.66×2.67 m). Each social group consisted of 6-7 male macaques of similar age, including both MS group and MR group monkeys and they lived for at least 6 months before the first behavioral sampling. Behavioral data were collected by video recordings when the monkeys were 1-1.5 years old.

### 2.2 Behavioral data collection and analysis

#### 2.2.1 Behavior definitions

According to the DSM-V and clinical diagnostic criteria for ASD, the core symptoms of ASD patients are impaired social interaction, stereotyped behaviors and restricted interest (Zhao, Jiang et al. 2018, Hodges, Fealko et al. 2020). Therefore, we focused on these three behaviors of macaques in this study.

Social interaction refers to all social interaction between the macaque and the other monkeys, including social contact, social approach, social grooming and clinging (Higley, Suomi et al. 1996, Martin, Ashwood et al. 2008). Social contact refers to huddling or sitting together in close contact and it includes two forms of interaction: active contact and passive contact by peers. Social approach means that the distance between macaques is less than one-arm length without contact, which including active approach and passive approach as well (Marais, Daniels et al. 2006). Grooming is an activity performed by macaques in order to keep their fur in good hygiene and neat appearance. Social grooming refers to grooming hair between macaques, usually in the form of stroking, scratching and massaging. It includes grooming others and being groomed by others (Higley, Suomi et al. 1996). Clinging is a special form of social interaction which commonly seen in infant monkeys, including ventral contact, tandem walking and draping an arm around a companion (Higley, Suomi et al. 1996). Restricted interest refers to overly restricted, persistent interest or intense fascination or preoccupation with one object, including interest in oneself and objects in the surroundings (Matson, Dempsey et al. 2009). Stereotyped behaviors refer to actions that is repeated three or more times with no apparent purpose and function, including various forms, such as digit-sucking, pacing, bouncing, body-flipping, rocking, and self-grasping. Among those behaviors, digit-sucking means sucking on a finger or toe. Pacing is a repetitive, ritualized movement that usually involves circling the cage. Bouncing is a monkey jumps up and down on all four legs. Body-flipping means grasping the top of the cage and flipping the body between the arms. Rocking is a back-and-forth movement of the upper body with stationary feet. Self-grasping refers to the monkey grabbing or holding onto a part of its own body (Mason 1991, Lutz, Well et al. 2003, Martin, Ashwood et al. 2008, Edwards, Lang et al. 2012).

#### 2.2.2 Video recording and behavioral data analysis

The collection and analysis of behavioral data were performed by three experienced observers. The entire procedure strictly adhered to the double-blind principle. To avoid sampling biases, each monkey was recorded both in the morning and afternoon at a specific time. Each monkey was collected 4-8 pieces of 30-minute videos (2-4 hours in total).

All monkeys had been familiar with the observers and cameras for a week before the behavioral video recording started. For video recording, the target macaque was first selected, and then a tripod, loaded with a digital camera, was mounted in front of the macaque’s living cage to record the macaque’s natural behaviors. To avoid interference with the monkeys, the observers and videotaped equipment were at least 5 m away from the cage. At the beginning of video recording, it was required to mark the characteristics of the target macaque as well as the recording date and time. During the video recording, when the loss of the target monkey occurred, it was corrected at once and the time of the loss was marked.

All behavioral data was analyzed on computers. Each video was used to analyze the behaviors of the targeted monkey. Three observers were required to analyze each video at the same time and reach an agreement at the end of each behavior. The stereotyped behaviors of monkey A are used below as an example to illustrate the specific steps of the behavioral analysis. When monkey A started any form of stereotyped behaviors (e.g., digit-sucking) in the video, manually pause the video and record the exact time point. Then continued to play the video until the stereotyped behavior stopped, pause again, and recorded the precise time point. The time interval between these two precise time points was the duration of this episode of stereotyped behavior for monkey A. In a video, there were usually multiple stereotyped behaviors. The sum of the durations of each stereotyped behavior was the duration (s) of monkey A’s stereotyped behaviors in this video, and the sum of the occurrence times of each stereotyped behavior was the frequency (f) of monkey A’s stereotyped behaviors in this video. Then we divided this duration (s) and frequency (f) of stereotyped behaviors by the total duration (h) of the video file to obtain the duration (s/h) and frequency (f/h) of stereotyped behaviors of the monkey A in unit time. Other behaviors were analyzed similarly.

### 2.3 Statistical analysis

Statistical analysis and plotting of all data were performed using SPSS V26.0 (SPSS Chicago, USA) and GraphPad Prism V9.0 (GraphPad Software, San Diego, USA). The analysis of differences between groups (Fig.1) was performed using the Mann Whitney U test. Differential analysis between one macaque and the control group (Fig.2 and Fig.3) was performed using one sample t test (the data satisfies normal distribution) and one sample Wilcoxon test (the data does not satisfy the normal distribution). Data are shown as mean ± SEM, \**p* < 0.05, \*\**p* < 0.01, \*\*\*\**p* < 0.0001. All *p* values were generated by two-tails tests.

**Fig.1.**
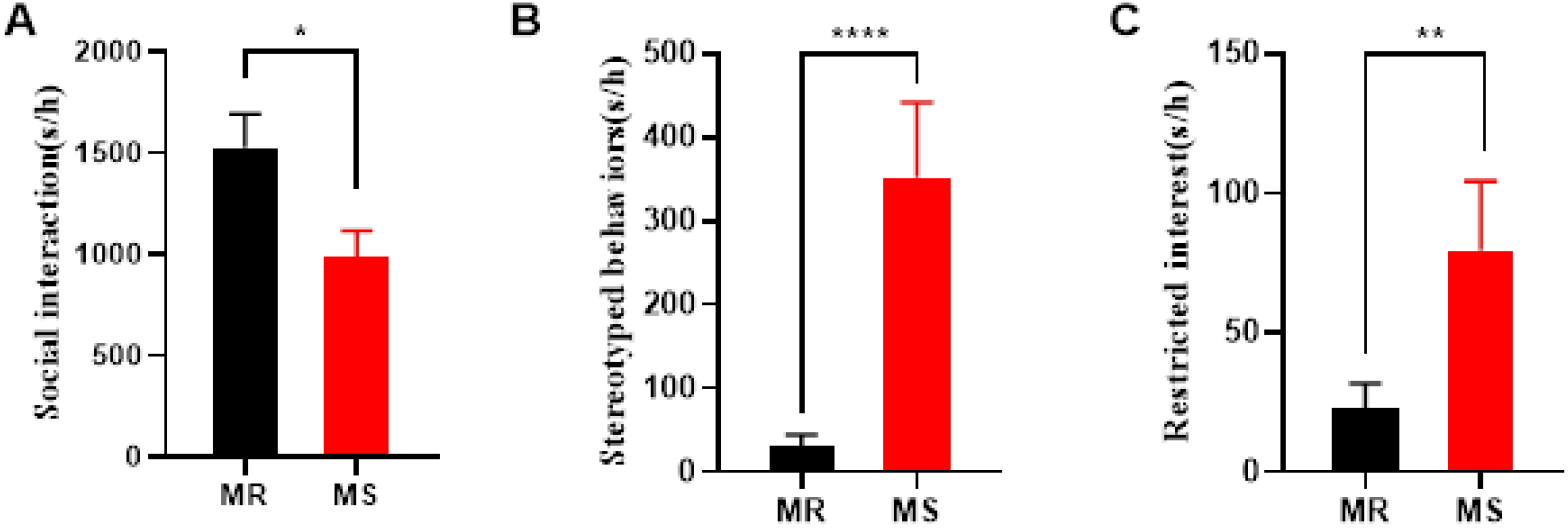
Comparison of the ASD core clinical symptoms between screened MS macaques (MS, male, 1-1.5 years, n=4) and MR controls of the same age (MR, male, 1-1.5 years, n=9). Compared with the MR group (black bar), the MS group (red bar) displayed (A) significant reduction in duration of social interaction (Mann Whitney test, U=305, \**p*=0.0170), (B) significantly increase in duration of stereotyped behaviors (Mann Whitney test, U=148.5, \*\*\*\**p*<0.0001) and (C) significantly increased duration of restricted interest (Mann Whitney test, U=299, \*\**p*=0.0030). Data are presented as mean ± SEM, * *p* < 0.05, ** *p* < 0.01, \*\*\*\**p* < 0.0001. All *p* values were generated by two-tails tests.

**Fig.2.**
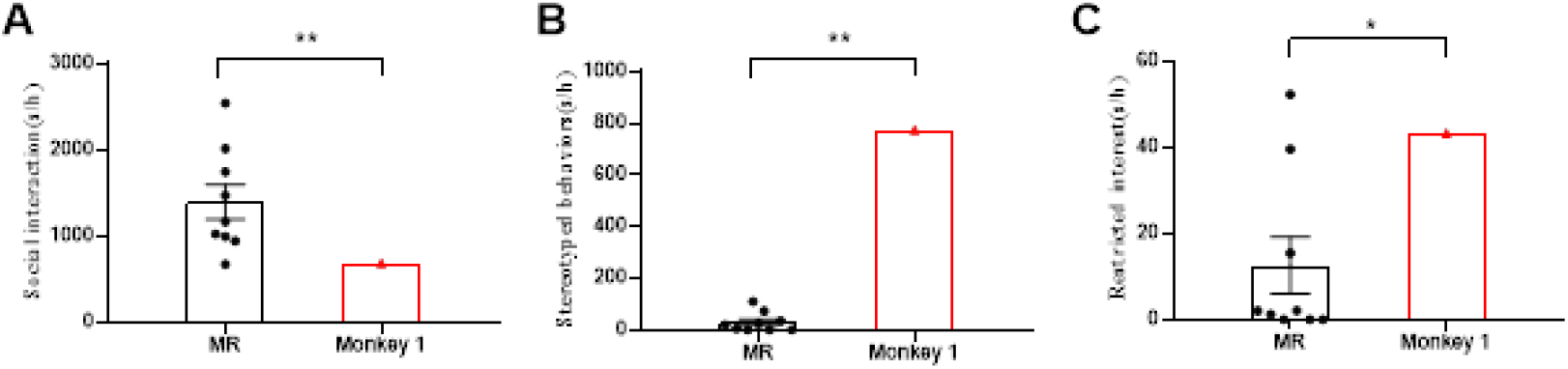
Comparison of the three ASD core clinical symptoms between Monkey1 and the age-matched MR controls (MR, male, 1-1.5 years old, n=9). Compared with the MR controls (black dots), Monkey 1 (red triangle) displayed (A) significant reduction in duration of social interaction (One sample t test, t=3.553, \*\**p*=0.0075), (B) significantly increased duration of stereotyped behaviors (One sample Wilcoxon test, \*\**p*=0.0039) and (C) significantly increased duration of restricted interest (One sample Wilcoxon test, \**p*= 0.0117). Data are presented as mean ± SEM, * *p* < 0.05, ** *p* < 0.01, \*\*\*\**p* < 0.0001. All *p* values were generated by two-tails tests.

**Fig.3.**
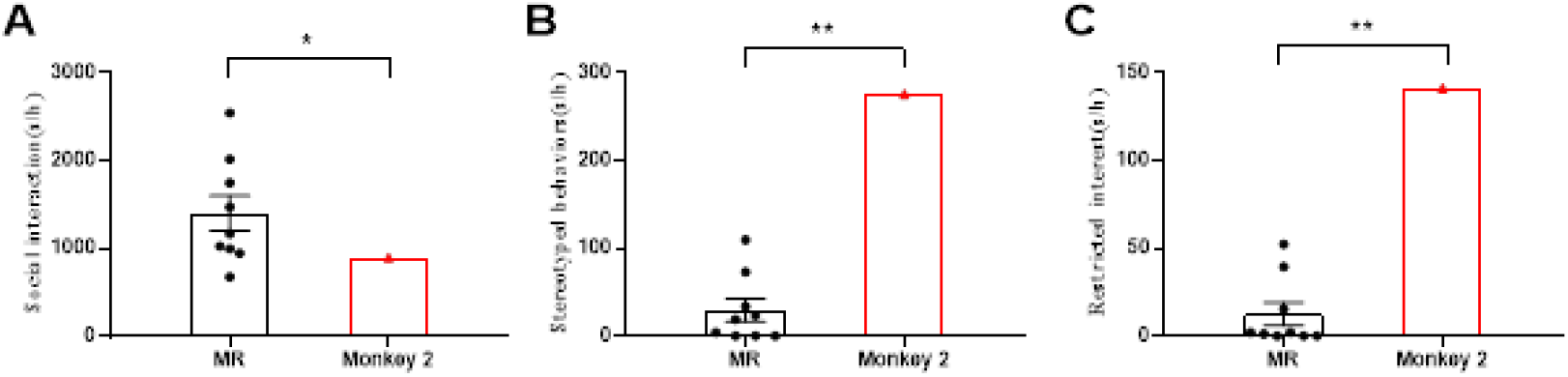
Comparison of the three ASD core clinical symptoms between Monkey2 and the age-matched MR controls (MR, male, 1-1.5 years old, n=9). Compared with the MR controls (black dots), Monkey 2 (red triangle) displayed (A) significant reduction in duration of social interaction (One sample t test, t=2.504, \**p*=0.0367), (B) highly increased duration of stereotyped behaviors (One sample Wilcoxon test, \*\**p*=0.0039) and (C) highly increased duration of restricted interest (One sample Wilcoxon test, \*\**p*=0.0039). Data are presented as mean ± SEM, * *p* < 0.05, ** *p* < 0.01, \*\*\*\**p* < 0.0001. All *p* values were generated by two-tails tests.

## 3. Results

To investigate whether maternal separation could induce ASD monkeys, we recorded and analyzed the behavioral data of 12 MS monkeys and four macaques with at least one of the three core clinical symptoms of ASD were identified. These four MS macaques as a group showed less social interaction, more stereotyped behaviors, and more restricted interest than the MR controls. This result suggested that there were possibly ASD individuals among the four monkeys. We further analyzed the individual behaviors of the four in detail and found two ASD young monkeys. Details of the results are presented below.

### 3.1 preliminary screening of candidate ASD monkeys

In the experiment, we first screened the young monkeys with ASD tendency in 12 MS macaques (male, 1-1.5 years old) that had undergone maternal separation. The behavioral data of the targeted monkey living in a mixed social group was collected by video, and then analyzed to investigate the differences of various ASD related behaviors between MS and MR monkey. According to DSM-V and clinical diagnostic criteria for ASD, impairments in social interaction, repetitive stereotyped behavior, and restricted interest are core clinical symptoms of ASD. Therefore, we assessed the 12 MS monkeys for social interaction, stereotyped behaviors, and restricted interest and found four monkeys among them with at least one of the three ASD core clinical symptoms. Then the four macaques were pulled together as a group (MS group, 1-1.5 years old, n=4) for the comparison with another same age group raised by their mothers (MR group, 1-1.5 years old, n=9). The results showed that the MS group displayed significantly reduced duration of social interaction (\**p*=0. 0107) (Fig.1 A), increased duration of stereotyped behaviors (\*\*\*\**p*<0.0001) (Fig.1 B) and increased duration of restricted interest (\**p*= 0.0140) (Fig.1 C). In other words, compared with the MR monkeys, the four MS monkeys overall showed significantly more ASD core clinical symptoms. We speculated that there were probably ASD monkeys that simultaneously had all the three symptoms among the four.

### 3.2 Identification of ASD monkeys

The results above suggested that there might be ASD monkeys among the four preliminarily screened candidate MS young monkeys. To identify them, we performed a detailed quantitative analysis on the behaviors of each one of the four, which was focused on the three core clinical symptoms of ASD, and compared them with the behaviors of the MR monkeys (male, 1-1.5 years old, n=9). It was found that two of the four monkeys exhibited all the core ASD symptoms. Compared with the nine MR macaques, Monkey 1 showed highly significant decrease in duration of social interaction (\*\**p*=0.0075) (Fig.2 A), and significant increase in duration of stereotyped behaviors (\*\**p*=0.0039) and restricted interest (\**p*=0.0117) (Fig.2 B, C). Similarly, Monkey 2 also displayed significantly more core symptoms of ASD including less social interaction (\**p*=0.0367), more stereotyped behaviors (\*\**p*=0.0039) and more restricted interest (\**p*=0.0039) than nine MR macaques (Fig.3 A, B, C).

Therefore, according to DSM-V and clinical criteria for ASD, we believe that Monkey 1 and Monkey 2 are two ASD macaques. These results demonstrate that, maternal separation, as an early environmental factor after birth, could induce all the three core clinical symptoms of ASD in some of young rhesus monkeys.

In addition to the two young monkeys with all the three core ASD clinical symptoms, the other two MS monkeys also exhibited partial core symptoms of ASD (Table. 1). Monkey 3 showed more stereotyped behaviors (One sample Wilcoxon test, \*\**p*=0.0078) and restricted interest (One sample Wilcoxon test, \*\**p*=0.0039) than the MR controls (1-1.5 years old, n=9). The duration of stereotyped behaviors (One sample t test, \**p*=0. 0296) was significantly higher in Monkey 4 than that of the MR controls.

**Table.1.** There are two monkeys (Monkey 1 and Monkey 2) exhibited all three core symptoms of ASD and the other two monkeys (Monkey 3 and Monkey 4) exhibited partial symptoms of ASD among the four MS monkeys screened from the twelve male macaques (1-1.5 years old) that underwent maternal separation.

| Monkey ID | Social<br>interaction | Stereotyped<br>behaviors | Restricted<br>interest |
| --- | --- | --- | --- |
| Monkey 1 | $**_{(a)}$ | $**_{(b)}$ | $^{*}_{(c)}$ |
| Monkey 2 | $^{*}_{(d)}$ | $**_{(e)}$ | $**_{(f)}$ |
| Monkey 3 | ns | $**_{(g)}$ | $**_{(h)}$ |
| Monkey 4 | ns | $**_{(i)}$ | ns |

ns: no significant difference; $**_{(a)}$ : One sample t test, $t=3.553$ , $**p=0.0075$ ; $**_{(b)}$ : One sample Wilcoxon test, $**p=0.0039$ ; $^{*}_{(c)}$ : One sample Wilcoxon test, $*p=0.0117$ ; $^{*}_{(d)}$ : One sample t test, $t=2.504$ , $*p=0.0367$ ; $**_{(e)}$ : One sample Wilcoxon test, $**p=0.0039$ ; $**_{(f)}$ : One sample Wilcoxon test, $**p=0.0039$ ; $**_{(g)}$ : One sample Wilcoxon test, $**p=0.0078$ ; $**_{(h)}$ : One sample Wilcoxon test, $**p=0.0039$ ; $**_{(i)}$ : One sample t test, $*p=0.0296$ .

## 4. Discussion

In order to explore whether maternal separation could induce core symptoms of ASD in rhesus monkeys, for the first time, we conducted a rigorous study on young macaques underwent maternal separation and age-matched normal macaques raised by their mothers. And two young MS monkeys were found to exhibit the three core symptoms of ASD, which are impairment in social interaction, abnormally increased stereotyped behaviors and restricted interest. According to DSM-V and ASD clinical diagnostic criteria, these two individuals should be ASD monkeys for having all the three core ASD symptoms simultaneously. There were also two young monkeys that showed partial symptoms of ASD, one showing abnormally increased in stereotyped behaviors and restricted interest, and the other showing only abnormal stereotyped behaviors. This is very similar to those of human beings. Clinical data demonstrated that many human ASD patients do not necessarily have significant deficits of all the three core symptoms of ASD at the same time, but rather exhibit only one or two of them.

The monkey ASD model induced by altering early environment in this study has similarities with the previous transgenic ASD monkey models and also has some differences in terms of clinical symptoms. For example, using a similar behavioral analysis protocol, a study we collaborated with Dr. Ji’s research team in 2017 showed that editing the MeCP2 gene via TALEN technology at embryonic stage resulted in two out of the three core symptoms: reduced social interaction and increased stereotyped behaviors in cynomolgus monkeys (Chen, Yu et al. 2017). In 2021, our group in collaboration with Dr. Qiu’s Lab used CRISPR/Cas9 *in situ* editing MeCP2 gene in the hippocampus of rhesus monkeys and induced only significant social impairment among the three symptoms (Wu, Li et al. 2021). In contrast to these two previous transgenic models that we took part in, this environmental factor ASD monkey model developed by our and Dr. Qiu’s Lab, showed all the three core symptoms of ASD at the same time and, in particular, showed significantly increased restricted interest for the first time. Therefore, this monkey model induced by early environmental factors was more comprehensive in terms of mimicking the core clinical symptoms of ASD.

This is the first report on maternal separation as an adverse environmental factor in early life can induce all the three core clinical symptoms of ASD in rhesus monkeys. These findings, on one hand, may provide a new approach for ASD mechanism study as well as early intervention and treatment of ASD; On the other hand, it provides technical support for establishing the disease model of ASD by using rhesus monkeys. An increasing number of studies have demonstrated that environmental factors are equally important compared to genetic factors in the pathogenesis of ASD. Therefore, it is of great importance to systematically investigate the environmental and genetic interactions in the pathogenesis of ASD. Combining this study with previous studies of editing ASD genes may provide a good foundation for future studies on the interaction between gene and environment in the development and progression of ASD.

## Author contributions

Xin-Tian Hu, Xiao-Li Feng and Zi-Long Qiu designed and led the project. Xiao-Feng Ren, Xiao-Li Feng and Hui Zhou performed the video recording and behavioral analysis. Xiao-Feng Ren and Shi-Hao Wu conducted statistical analysis of the data. Xiao-Feng Ren and Xin-Tian Hu prepared the manuscript. Xiao-Li Feng, Shi-Hao Wu and Zi-Long Qiu played critical roles in the revision of the article. All authors read and approved the final version of the manuscript.

## Acknowledgement

This study was supported by the Key-Area Research and Development Program of Guangdong Province (2019B030335001), National Key Research and Development Program of China (2021YFF0702700, 2018YFA0801403), Strategic Priority Research Program of the Chinese Academy of Sciences (CAS) (XDB32060200), Science and Technology Service Network Initiative (STS) Project of Chinese Academy of Sciences (E02E1801), National Natural Science Foundation of China (81941014, 81771387, 31800901, 31960178, 31700910), Applied Basic Research Programs of Science and Technology Commission Foundation of Yunnan Province (202001AT070130, 202101AT070437, 202101AT070403), Kunming Medical Joint Project-Key Project (202101AY070001-001)), Yunnan Province Ten Thousand Talents Program Young Top Talent Special Project (YUWR-QNBJ-2019-043), CAS “Light of West China” Program, and Kunming Medical University 100 young and middle-aged academic and technical backbone training program (J13326055). We would like to thank the National Research Facility for Phenotypic and Genetic Analysis of Model Animals, Kunming Institute of Zoology, Chinese Academy of Sciences and the National Resource Center for Non-Human Primates for technical support and animal care.

